# Discovery of dihydroxy-enone–type protein-bound ceramides as the dominant type in human stratum corneum

**DOI:** 10.64898/2026.04.08.717327

**Authors:** Ayumi Kojima, Takumi Sugiyama, Yusuke Ohno, Akio Kihara

**Affiliations:** Faculty of Pharmaceutical Sciences, Hokkaido University, Sapporo 060-0812, Japan

**Keywords:** ceramide, epidermis, lipid, protein-bound ceramide, skin barrier

## Abstract

Protein-bound ceramides are a specialized subclass of ceramides that are essential for skin barrier function, and their defective formation leads to severe skin disorder ichthyosis. Despite their biological importance, the precise molecular structures of protein-bound ceramides have remained incompletely defined, largely due to the technical challenges arising from their unique covalent linkage between lipid and protein components with highly distinct physicochemical properties. Using mass-spectrometry–based analyses of acylceramide moieties derived from protein-bound ceramides, we investigated whether epoxy-enone (EE)–type protein-bound ceramides present in mouse epidermis are conserved in human skin. Although EE-type protein-bound ceramides were detectable in the human stratum corneum, they constituted only a minor fraction of the ceramides in this tissue. Instead, we discovered a previously unrecognized class of protein-bound ceramides, termed dihydroxy-enone (DE)–type protein-bound ceramides, as the predominant class in human skin. These DE-type ceramides are generated through hydrolytic opening of the epoxide moiety of EE-type ceramides. In contrast, DE-type protein-bound ceramides were present in mouse epidermis at much lower levels. DE-type acylceramides appeared as two chromatographically distinct peaks, which likely correspond to putative stereoisomers with (9*R*,10*S*) and (9*R*,10*R*) configurations. Age-dependent increases in the (9*R*,10*S*) form in mouse epidermis closely paralleled changes in the expression levels of the epoxide hydrolase *Ephx3*, suggesting a role for EPHX3 in the conversion of EE- to DE-type ceramides. Together, these findings further reveal molecular diversity in protein-bound ceramides and a fundamental difference between human and mouse epidermal lipid architectural organization.

## Introduction

The stratum corneum (SC), the outermost layer of the epidermis, functions as a skin barrier (permeability barrier) that reduces transepidermal water loss and prevents the invasion of external substances such as pathogens, allergens, and chemicals. The SC is composed of corneocytes (terminally differentiated dead keratinocytes) and intercellular multilayered lipid structures (lipid lamellae) that fill the spaces between the corneocytes (1, 2). Ceramides are the predominant lipid components of the lipid lamellae and play a critical role in skin barrier function, together with cholesterol and free fatty acids (3, 4).

Ceramides are composed of a long-chain base and a fatty acid (*N*-acyl chain), which are linked via an amide bond (Fig. 1*A*). Ceramides are classified into two major categories: free (non–protein-bound) ceramides and protein-bound ceramides. Free ceramides are structural components of the lipid lamellae, whereas protein-bound ceramides are constituents of the lipid structure covering corneocytes, known as the corneocyte lipid envelope (CLE) (5–7). Unlike other cells, which possess a phospholipid bilayer plasma membrane, corneocytes have instead a highly robust CLE, which provides a scaffold for the extracellular lipid lamellae. Protein-bound ceramides have an *N*-acyl chain covalently attached to corneocyte surface proteins (the cornified envelope proteins). Free ceramides are further classified into non-acylated ceramides (the conventional type) and ω-*O*-acylceramides (hereafter referred to as acylceramides) (3, 4). Acylceramides possess an *N*-acyl chain that is ω-hydroxylated, and this ω-hydroxyl group is esterified with an additional *O*-acyl chain (predominantly linoleic acid) (Fig. 1*A*). Non-acylated ceramides serve as the fundamental building blocks of the lipid lamellae, whereas acylceramides, although less abundant, play a crucial role in the formation and maintenance of the lipid lamellae (4, 8–11). Among ceramides, acylceramides and protein-bound ceramides are particularly important for skin barrier function, since mutations in the genes responsible for their synthesis cause the severe skin disorder ichthyosis (also known as an epidermal differentiation disorder) in humans (12, 13), and knockout (KO) of the corresponding genes in mice results in neonatal lethality due to severe skin barrier defects (9, 14–25).

**Fig. 1.**
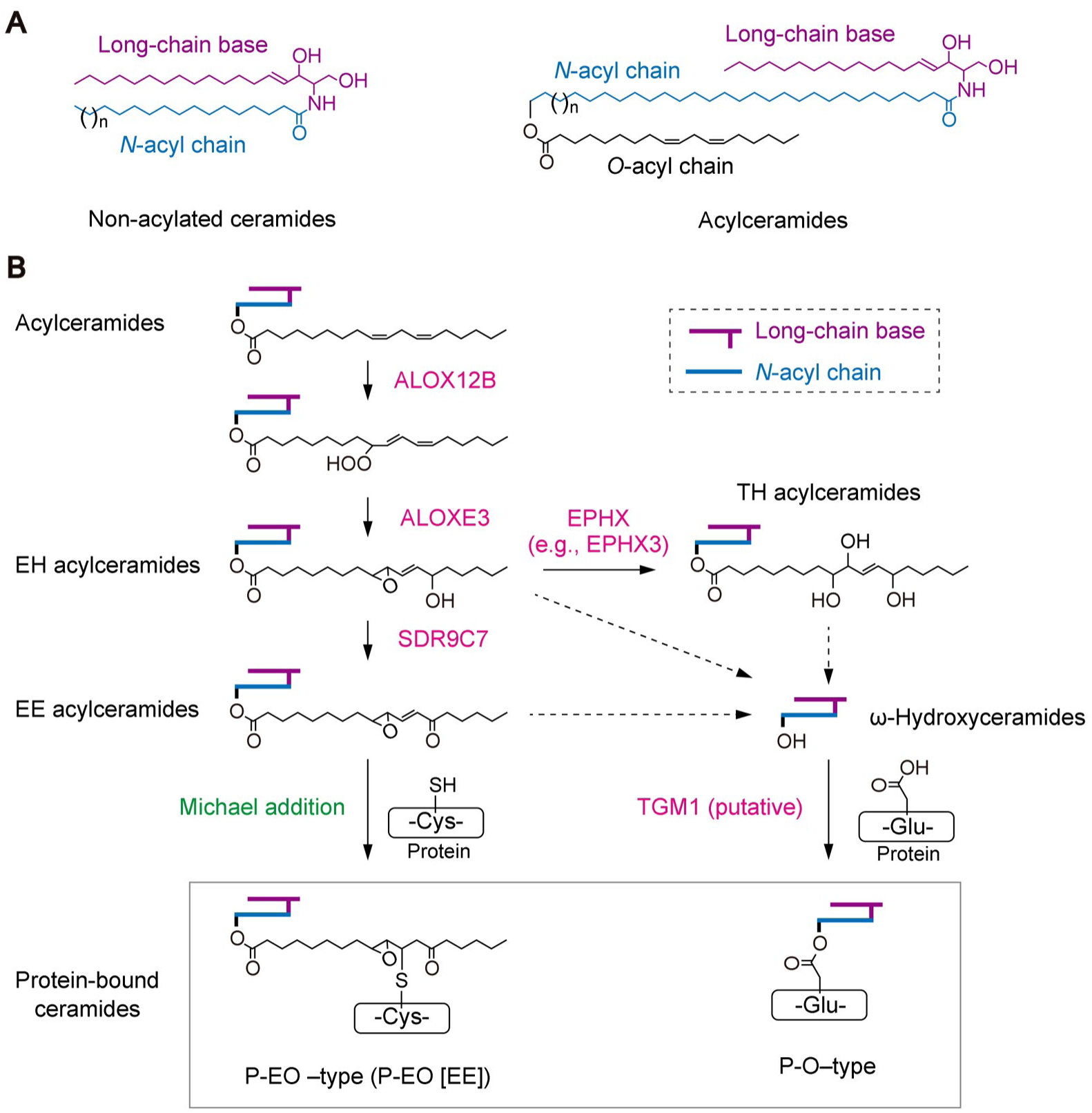
Ceramide structures and proposed biosynthetic pathways for P-EO–type and P-O–type protein-bound ceramides. (*A*) Structures of non-acylated ceramides and acylceramides. (*B*) Proposed biosynthetic pathways for P-EO–type and P-O–type protein-bound ceramides. The linoleic acid moiety of acylceramides undergoes hydroperoxidation catalyzed by ALOX12B, followed by hydroperoxide isomerization by ALOXE3, resulting in the formation of EH acylceramides. EH acylceramides are then converted into EE acylceramides or TH acylceramides by SDR9C7 or epoxide hydrolases, including EPHX3, respectively. EE acylceramides covalently bind to cysteine residues of corneocyte surface proteins via Michael addition, yielding P-EO–type protein-bound ceramides (P-EO [EE]). In the proposed pathway for the formation of P-O–type protein-bound ceramides, the *O*-acyl chains of EH, EE, and/or TH acylceramides are cleaved by an as-yet-unidentified esterase to yield ω-hydroxyceramides, which are subsequently covalently bound to glutamate residues of corneocyte surface proteins. Although TGM1 has been considered as a candidate enzyme, its involvement in this process remains debated. Dashed arrows indicate putative pathways for which the responsible enzymes and/or reaction mechanisms remain unknown.

Two structural models have been proposed for protein-bound ceramides (Fig. 1*B*) (23, 26). One is the P-O model, in which the fatty acid (*N*-acyl) moiety of a protein-bound ceramide is represented by the P-O unit, which is an abbreviation for <u>p</u>rotein-bound <u>ω</u>-hydroxy fatty acid. P-O–type ceramides are designated, for example, as P-OS, where P-O represents the *N*-acyl chain and S represents the long-chain base (abbreviated from <u>s</u>phingosine). In this model, the ω-hydroxyl group of the ω-hydroxy fatty acid moiety is ester linked to a protein. The other structural model is the P-EO model, in which the *N*-acyl moiety of a protein-bound ceramide consists of an ω-hydroxy fatty acid esterified with a modified linoleic acid; P-EO designates a <u>p</u>rotein-bound modified linoleic acid–<u>e</u>sterified <u>ω</u>-hydroxy fatty acid (23, 27). In this model, the modified linoleic acid moiety is covalently bound to a protein.

The P-O model has long been accepted as the structural model for protein-bound ceramides, but there have been no studies published that directly demonstrate the existence of these forms at the molecular level. However, we recently revealed the existence of P-EO–type protein-bound ceramides, which originate from epoxy-enone (EE) acylceramides (27). These EE acylceramides are generated by sequential hydroperoxidation, isomerization of the resulting hydroperoxide, and oxidation of the linoleic acid moiety of acylceramides mediated by the lipoxygenases ALOX12B and ALOXE3 and the short-chain dehydrogenase/reductase SDR9C7 (Fig. 1*B*) (23, 28). Since the enone group is highly reactive in general, it has been suggested that EE acylceramides may non-enzymatically form covalent bonds with corneocyte surface proteins (23). Indeed, we showed *in vitro* that the β-carbon of the enone moiety of EE acylceramides binds to the thiol group of cysteine residues via a Michael addition reaction (27). Furthermore, we detected EE acylceramide–cysteine conjugates in mouse epidermis *in vivo* after proteolytic digestion of the protein moiety of protein-bound ceramides into amino acids using Pronase E. However, we did not detect P-O–type protein-bound ceramides in our analyses.

The biosynthetic pathway proposed for P-O–type ceramides is as follows: the modified linoleic acid moiety of acylceramides (EE acylceramides, their precursors epoxyhydroxy [EH] acylceramides, and/or trihydroxy [TH] acylceramides generated by epoxide ring opening of EH acylceramides) is hydrolyzed to ω-hydroxyceramides by an as-yet-unidentified esterase (Fig. 1). The ω-hydroxyl group then reacts with glutamate residues of corneocyte surface proteins to form a covalent bond, generating P-O–type protein-bound ceramides. At least *in vitro*, this reaction can be catalyzed by the transglutaminase TGM1 (29). However, it remains controversial whether TGM1 catalyzes this reaction *in vivo*, or even whether this reaction actually occurs *in vivo* (10, 30).

The composition of free ceramides in the SC differs markedly between humans and mice. For example, humans possess higher levels of phytosphingosine and 6-hydroxysphingosine as long-chain base moieties, while mice lack 6-hydroxysphingosine, contain lower levels of phytosphingosine, and instead have higher levels of sphingosine (31, 32). The two species also differ in the hydroxylation sites of the *N*-acyl moieties: humans have higher levels of α-hydroxy fatty acids, while mice predominantly contain β- or ω-hydroxy fatty acids (31). These interspecies differences in free ceramide composition led us to consider that the structure of protein-bound ceramides might also differ between humans and mice. In this study, we identified previously unrecognized P-EO–type protein-bound ceramides in the human SC. These ceramides are generated by conversion of the linoleate moiety from an EE to a dihydroxy-enone (DE) structure and are hereafter referred to as P-EO (DE) ceramides. P-EO (DE) ceramides were abundant in the human SC, whereas EE-containing P-EO–type ceramides (P-EO [EE]), which are predominant in mice (27), were present only at low levels. Furthermore, we obtained insights into the stereochemical properties of P-EO (DE) ceramides and the enzymes involved in their formation. Collectively, our findings provide a molecular basis for understanding the role of protein-bound ceramides in skin barrier formation.

## Results

### Identification of DE-containing P-EO–type protein-bound ceramides in human SC

We previously identified P-EO (EE) ceramides as the predominant protein-bound ceramides in mouse epidermis (27). In these ceramides, the enone group of the EE-modified linoleic acid moiety undergoes a Michael addition with cysteine residues of corneocyte surface proteins. However, it remained unclear whether such P-EO (EE) ceramides also constituted the principal human protein-bound ceramides. The ceramide moiety of P-EO (EE) ceramides (i.e., EE acylceramides) can be released by oxidative sulfoxide elimination upon incubating protein-bound ceramide fractions (the residual materials of epidermis or SC after removal of free lipids) in organic solvents (27, 28). In this study, exploiting this property, we investigated the presence of P-EO (EE) ceramides in human SC collected by tape stripping. The ceramides thus released were analyzed via liquid chromatography (LC)–tandem mass spectrometry (MS/MS) using a precursor-ion scan analysis in the positive-ion mode (third quadrupole [Q3], *m/z* = 264, corresponding to the C18:1 sphingosine structure; scan range, *m/z* 900–1200), which detects sphingosine-containing ceramides. In the mouse epidermis that was used as a control, major ion peaks were detected at *m/z* 1,051/1,069 and 1,079/1,097 corresponding to EE acylceramides containing C32:1 and C34:1 *N*-acyl chains (ω-hydroxy fatty acids), respectively (Fig. 2*A*). As sphingosine-based ceramides readily undergo dehydration during ionization (33), both [M+H]^+^ and [M+H−H_2_O]^+^ ions, which differ by 18 Da, appear at the same retention time. However, these EE acylceramide-related ions were barely detected in human SC. Instead, a pair of prominent ion peaks not assigned to EE acylceramides appeared at earlier retention times, at 12.8 min (peak 1) and 13.1 min (peak 2). A second pair of prominent ion peaks was detected at 13.5 min (peak 3) and 13.6 min (peak 4). The first ion peak pair (peaks 1 and 2) primarily contained two common precursor ions at *m/z* 1,043 and 1,061 (Fig. 2*B*), whereas the second pair (peaks 3 and 4) contained ions at *m/z* 1,071 and 1,089 (Fig. 2*A*). The ions at *m/z* 1,043 and 1,071 are presumed to represent the dehydrated forms of the ions at *m/z* 1,061 and 1,089, respectively, because each ion pair differs by 18 Da. Although each ion peak pair appeared as two distinct chromatographic peaks, the peaks in each pair had identical *m/z* values, indicating that they represented stereoisomers.

**Fig. 2.**
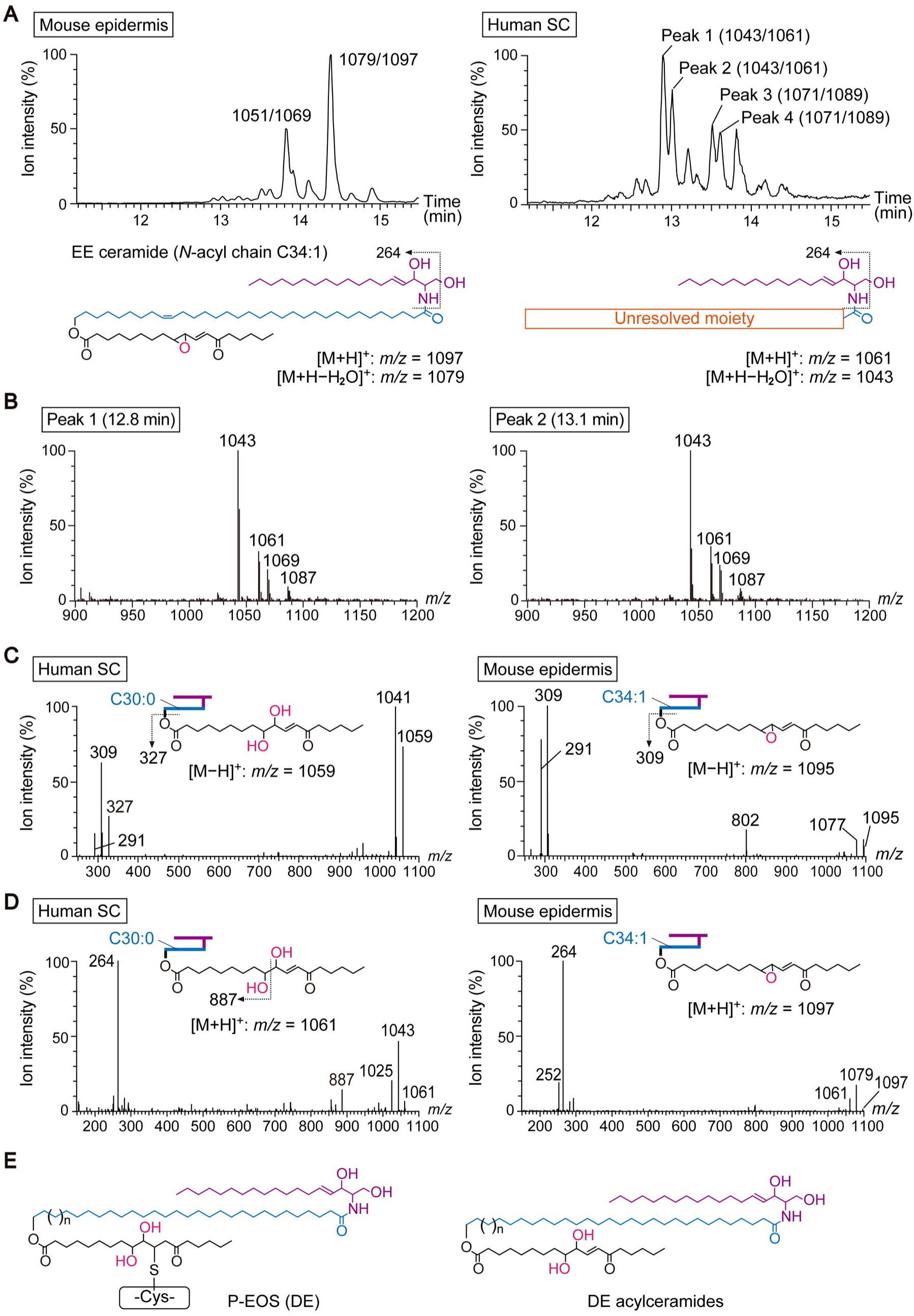
Identification of DE-containing P-EO–type protein-bound ceramides in human SC. (*A*–*D*) Protein-bound ceramide fractions were prepared from human SC and P0 mouse epidermis, and modified acylceramide moieties were released from P-EO–type protein-bound ceramides by oxidative sulfoxide elimination, followed by LC–MS/MS analysis. Total ion chromatograms (*A*) and mass spectra (*B*–*D*), together with the corresponding ceramide structures and their characteristic ions (*A*, *C*, and *D*), are shown. (*A* and *B*) Precursor ion scanning analyses (Q3, *m/z* 264; scan range, *m/z* 900–1200; positive ion mode) were performed to detect ceramides with a sphingosine backbone, shown as total ion chromatograms of mouse epidermis and human SC (*A*) and the corresponding mass spectra for peaks 1 and 2 detected in the human SC chromatogram (*B*). (*C* and *D*) Product ion scanning analyses were performed in the negative (*C*; Q1, *m/z* 1,059 [human SC] and *m/z* 1,095 [mouse epidermis]; scan range, *m/z* 250–1,100) and positive ion modes (*D*; Q1, *m/z* 1,061 [human SC] and *m/z* 1,097 [mouse epidermis]; scan range, *m/z* 250–1,100). In the schematic diagrams, the long-chain base and *N*-acyl chain are shown in purple and blue, respectively. (*E*) Structures of protein-bound ceramides (P-EO [DE] ceramides) identified in human SC and of DE acylceramides released by oxidative sulfoxide elimination. The structures of P-EOS (DE) and EOS (DE) are shown; these represent the sphingosine-containing forms of P-EO [DE] ceramides and DE acylceramides, respectively. Ceramide classes are represented by combinations of abbreviations for the *N*-acyl chain and long-chain base moieties. EOS indicates ceramides composed of the *N*-acyl chain moiety EO (<u>e</u>sterified <u>ω</u>-hydroxy fatty acid) and the long-chain base moiety S (<u>s</u>phingosine).

To further characterize the structure of these molecular ions, we performed product ion scanning analysis in the negative ion mode. The deprotonated ion ([M−H]^−^; *m/z* 1,059), corresponding to the protonated ion ([M+H]^+^; *m/z* = 1,061) that was detected with the highest ion intensity in the precursor ion scanning analysis (Fig. 2*A*, peaks 1 and 2), was selected as the precursor ion. In human SC, the precursor ion at *m/z* 1,059, its dehydrated form at *m/z* 1,041, and its characteristic product ions at *m/z* 309 and 327 were detected (Fig. 2*C*). In mouse epidermis, the deprotonated ion at *m/z* 1,095 ([M−H]^−^), corresponding to the most abundant EE acylceramide species with a C34:1 *N*-acyl chain (Fig. 2*A*), was likewise selected as the precursor ion. The precursor ion at *m/z* 1,095 and its dehydrated form at *m/z* 1,077 were detected, together with product ions at *m/z* 309 and 291 (the dehydrated form of *m/z* 309) from this EE acylceramide (Fig. 2*C*). In human SC, earlier elution on reversed-phase LC relative to EE acylceramides, along with the detection of a product ion at *m/z* 327, which is 18 Da higher than the corresponding EE acylceramide ion (*m/z* 309), suggests that the acylceramide is a hydrated form of EE acylceramide (i.e., DE acylceramide) in which the epoxy moiety is hydrolyzed to form a diol structure.

Following product ion scanning analysis in the negative ion mode (Fig. 2*C*), the same molecular species with a C30:0 *N*-acyl chain detected in human SC was further analyzed by product ion scanning analysis in the positive ion mode, using the protonated ion at *m/z* 1,061 ([M+H]^+^) as the precursor ion, to obtain additional structural information. In addition to the diagnostic product ion at *m/z* 264, which is derived from the sphingosine backbone, a product ion at *m/z* 887 was detected in human SC (Fig. 2*D*). In contrast, the product ion spectra obtained from the predominant EE acylceramide species, which has a C34:1 *N*-acyl chain, in mouse epidermis did not produce a fragment ion at *m/z* 923 corresponding to the *m/z* 887 fragment observed for the C30:0 species in human SC. The product ion at *m/z* 887 observed in human SC is assumed to arise from cleavage of the diol moiety, which supports the structural assignment of the detected species as a DE acylceramide. Collectively, these results indicate that DE acylceramides are released from human SC by oxidative sulfoxide elimination, revealing the presence of previously unrecognized P-EO (DE)–type protein-bound ceramides in human SC (Fig. 2*E*).

### Differential abundance of P-EO (EE) and P-EO (DE) ceramides in humans and mice

To quantify P-EO (EE) and P-EO (DE) ceramides in humans and mice, the corresponding acylceramides (EE and DE acylceramides, respectively) released from protein-bound ceramide fractions by oxidative sulfoxide elimination were quantified via LC–MS/MS using the multiple reaction monitoring (MRM) mode. MRM enables highly specific and sensitive detection of target ceramide species by monitoring defined precursor ions in the first quadrupole (Q1), corresponding to ceramide species with different *N*-acyl moieties, and the sphingosine-derived product ion at *m/z* 264 in Q3. Although EE acylceramides were not detected in human SC by precursor ion scanning analysis because of this method’s lower sensitivity (Fig. 2*A*), MRM analysis revealed the presence of both EE and DE acylceramides (Fig. 3*A*). In human SC, the quantities of DE acylceramides were approximately six-fold those of EE acylceramides. Both ceramide types showed similar *N*-acyl chain compositions, with C30:0 being the most abundant (Fig. 3*B*). In contrast, in the epidermis of postnatal day 0 (P0) mice, only trace quantities of DE acylceramides were detected, comprising approximately 7% of EE acylceramide quantities (Fig. 3*C*). In mouse epidermis, C34:1 was the predominant *N*-acyl chain species for both EE and DE acylceramides (Fig. 3*D*). The DE/EE acylceramide ratio was markedly higher in human SC than in mouse epidermis (Fig. 3*E*). Taken together, these quantitative analyses show that the predominant type of protein-bound ceramides differs markedly between humans and mice.

**Fig. 3.**
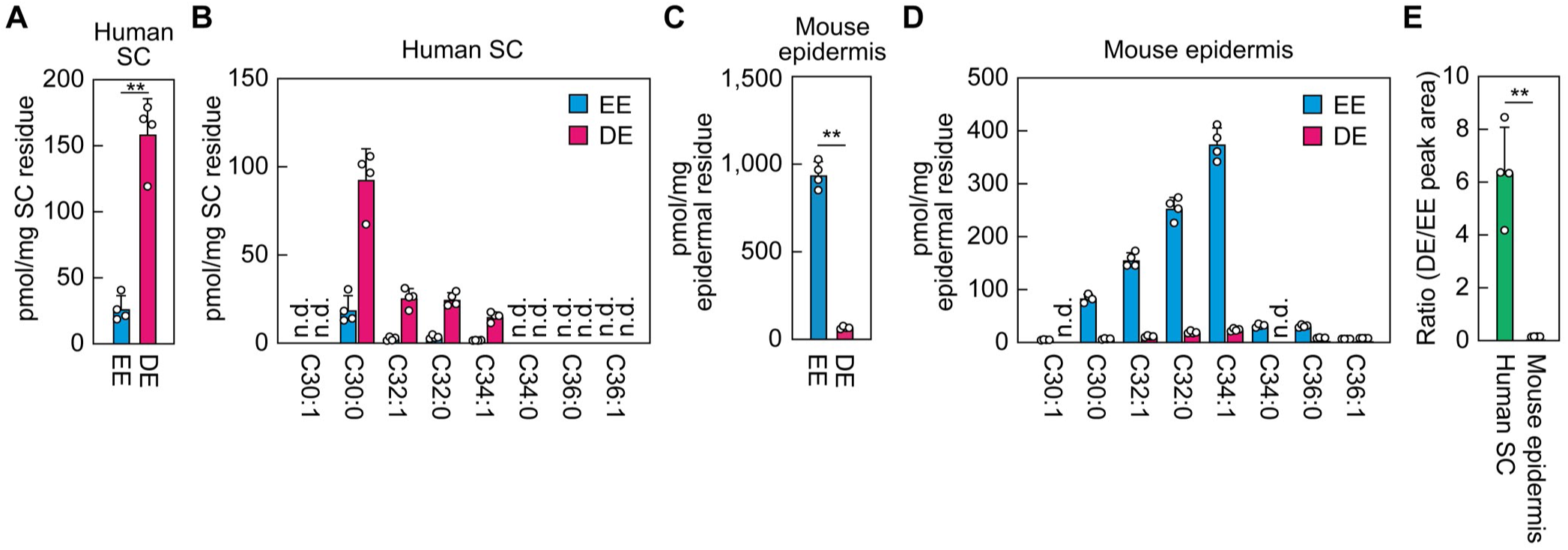
Differential abundance of P-EO (DE) and P-EO (EE) ceramides in human SC and mouse epidermis. Protein-bound ceramide fractions were prepared from human SC (*A* and *B*) and P0 mouse epidermis (*C* and *D*). Following oxidative sulfoxide elimination, modified acylceramide moieties were released from P-EO–type ceramides and quantified via LC–MS/MS in the MRM mode. The total quantities of DE and EE acylceramides (*A* and *C*), quantities of the individual DE or EE acylceramide species categorized by *N*-acyl chain moiety (*B* and *D*), and the ratio of DE acylceramides relative to EE acylceramides (*E*) are shown. Values presented are means + SD (n = 4; ** *P* < 0.01; Welch’s *t*-test). n.d., not detected.

### Predominance of reversibly releasable protein-bound ceramides in human SC

Based on their releasability via oxidative sulfoxide elimination, protein-bound ceramides can be divided into reversible and irreversible types (Fig. 4*A*). P-EO (EE) and P-EO (DE) ceramides are classified as the reversible type, because their corresponding acylceramides can be released by this reaction. We previously showed that approximately 60% of protein-bound ceramides in mouse epidermis are the reversible type, most of which are P-EO (EE) ceramides, while the remaining protein-bound ceramides are the irreversible type (27). Although the structural details of the irreversible type remain unclear, they may be P-O ceramides or P-EO ceramides that are not P-EO (EE) or P-EO (DE) ceramides. Regardless of reversibility, all protein-bound ceramides can be converted to ω-hydroxyceramides upon alkaline hydrolysis of the ester bond at the ω-position of the *N*-acyl chain (Fig. 4*B*). Thus, quantitative comparison of reversible and irreversible protein-bound ceramides is possible after conversion to ω-hydroxyceramides by alkaline treatment.

**Fig. 4.**
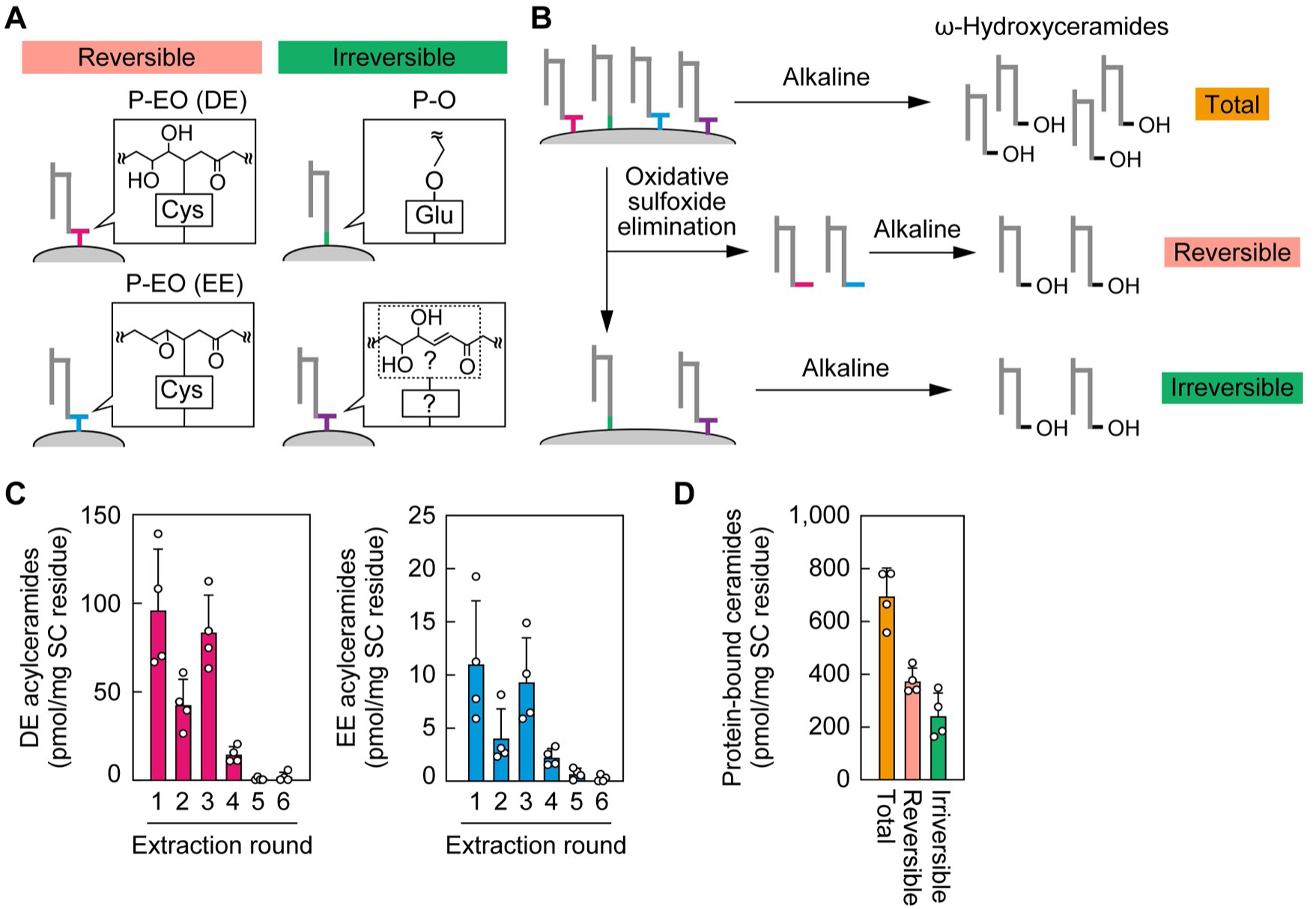
Abundance of reversibly and irreversibly bound protein-bound ceramides in human SC. (*A*) Schematic representation of both reversibly bound protein-bound ceramides that are released via oxidative sulfoxide elimination upon incubation of the protein-bound ceramide fraction in organic solvent and irreversibly bound protein-bound ceramides that remain unreleased. P-EO (EE) and P-EO (DE) ceramides are classified as reversibly bound protein-bound ceramides, whereas P-O ceramides are classified as irreversibly bound protein-bound ceramides. The irreversibly bound fraction may also include protein-bound ceramides derived from P-EO–type ceramides, although their structures have not yet been elucidated. (*B*) Schematic overview of the preparation and quantification of total, reversible, and irreversible fractions of protein-bound ceramides. The protein-bound ceramide fraction without treatment (total), the fraction released by oxidative sulfoxide elimination (reversible fraction), and the residual unreleased fraction (irreversible fraction) were each subjected to alkaline treatment to convert all types of protein-bound ceramides to ω-hydroxyceramides, which were then quantified via LC–MS/MS. (*C* and *D*) Protein-bound ceramide fractions were prepared from human SC, suspended in chloroform/methanol (1:2, v/v), and incubated at 60 °C to induce oxidative sulfoxide elimination. The incubation was performed six times (15 h for the third incubation and 3 h for the others). (*C*) Quantities of DE and EE acylceramides released in each eluate were quantified via LC–MS/MS in the MRM mode. Values presented are means + SD (n = 4). (*D*) The protein-bound ceramide fraction (total), the combined eluates from six incubations (reversible fraction), and the pellet remaining after the sixth incubation (irreversible fraction) were each subjected to alkaline treatment to convert all types of protein-bound ceramides to ω-hydroxyceramides, followed by quantification via LC–MS/MS in the MRM mode. Values presented are means + SD (n = 4).

Protein-bound ceramides in human SC were fractionated according to their releasability by oxidative sulfoxide elimination. The reversible type was defined as protein-bound ceramides released during six consecutive rounds of elution with CHCl_3_/CH_3_OH (1:2, v/v; fractions 1–6), whereas the irreversible type corresponded to the residual fraction remaining after this treatment. Both DE and EE acylceramides were detected mainly in fractions 1–4 (Fig. 4*C*), whereas only trace quantities were detected in fractions 5 and 6, indicating that reversibly releasable protein-bound ceramides were almost completely released over the course of the elution procedure. In addition to the reversible and irreversible fractions, total protein-bound ceramides were prepared from the protein-bound ceramide fraction without prior organic solvent treatment. Each of these fractions was subsequently subjected to alkaline treatment to liberate ω-hydroxyceramides, which were quantified via LC–MS/MS. Quantitative analysis revealed that reversible- and irreversible-type protein-bound ceramides accounted for approximately 60% and 40% of the total protein-bound ceramides in human SC, respectively (Fig. 4*D*). Thus, the proportion of reversible protein-bound ceramides relative to the total was similar between humans (Fig. 4*D*) and mice (27), although the predominant forms differed, being P-EO (DE) ceramides in human SC and P-EO (EE) ceramides in mouse epidermis (Fig. 3*E*).

### Abundance of P-EO (DE) and P-EO (EE) ceramides in mouse skin: effects of tissue compartment and postnatal stages

Quantities of P-EO (DE) and P-EO (EE) ceramides differed between human SC and mouse epidermis (Fig. 3). This variation may be attributable to differences in the sampling layers, since SC samples obtained by tape stripping represent only the upper layers of the SC, whereas epidermal samples encompass the entire SC. Although epidermal samples include other epidermal layers (e.g., the stratum basale, stratum spinosum, and stratum granulosum), protein-bound ceramides are predominantly localized in the SC. Therefore, the levels of protein-bound ceramides measured in the epidermis largely reflect those present in the SC. A further potential complication is that differences in age at sampling (adult humans versus neonatal mice) may also have contributed to the observed differences in abundance of P-EO (DE) and P-EO (EE) ceramides.

First, to examine whether the difference in DE/EE abundance was attributable to the sampling layer, the full epidermis or tape-stripped SC was prepared from P0 mice. After removal of free lipids, DE and EE acylceramides were released from P-EO (DE) and P-EO (EE) ceramides present in the residual material by oxidative sulfoxide elimination and quantified via LC–MS/MS. The levels of EE acylceramides were approximately 10-fold those of DE acylceramides in the epidermis and approximately five-fold in the tape-stripped SC samples (Fig. 5*A*). The DE/EE ratio in the SC was thus double that in the epidermis (0.22 vs. 0.11; Fig. 5*B*). This finding indicates that the DE/EE ratio differs between the upper layers of the SC and the SC as a whole, which suggests that conversion from EE to DE ceramides progresses during SC maturation.

**Fig. 5.**
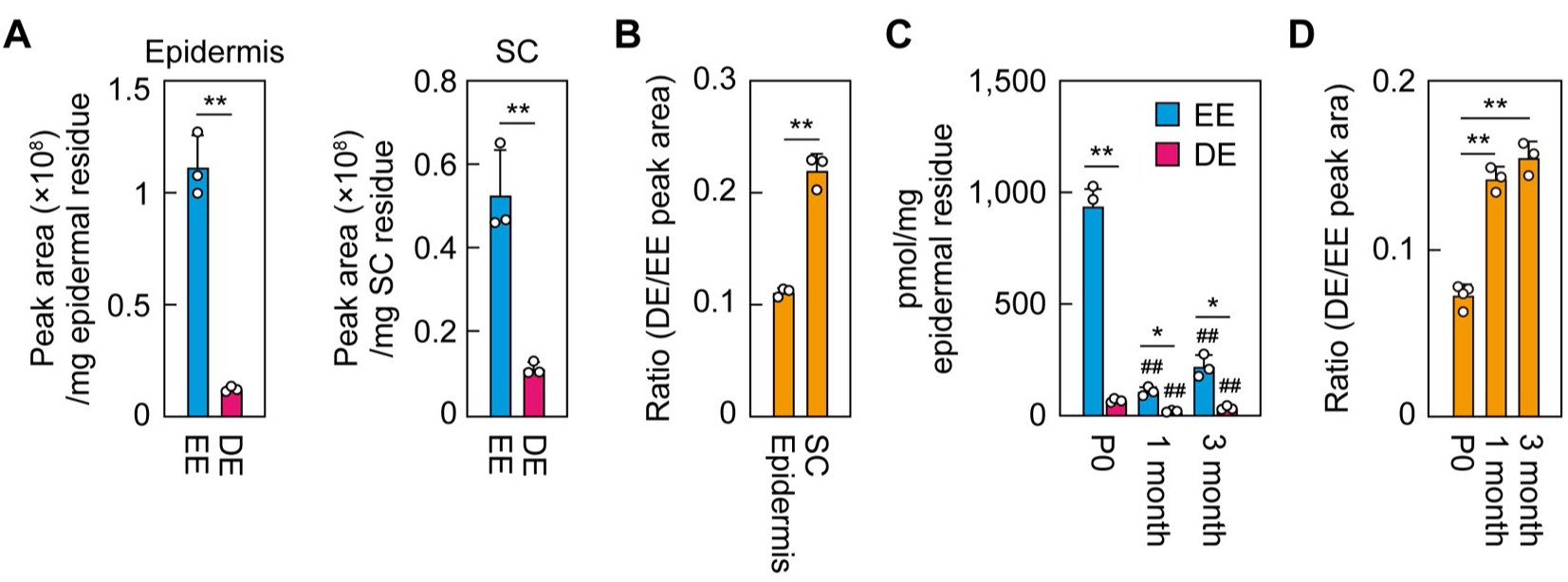
Regional and developmental variation of P-EO (DE) and P-EO (EE) ceramides in mouse skin. Protein-bound ceramide fractions were prepared from mouse epidermis or from SC, and DE and EE acylceramides were released via oxidative sulfoxide elimination and quantified via LC–MS/MS in the MRM mode. (*A* and *B*) Quantities of EE and DE acylceramides (*A*) and the ratio of DE to EE acylceramides (*B*) in epidermis and SC from P0 mice are shown. (*C* and *D*) Quantities of EE and DE acylceramides (*C*) and the ratio of DE to EE acylceramides (*D*) in the epidermis of P0 and one- and three-month-old mice are shown. Values presented are means + SD (n = 3; * *P* < 0.05, ** *P* < 0.01; Welch’s *t*-test [*A*–*C*], Tukey’s test [*D*]; *^##^ P* < 0.01; Dunnett’s test vs P0 mice [*C*]).

Next, to examine whether the difference in the DE/EE ratio was attributable to age, we quantified DE and EE acylceramides prepared from the epidermis of P0, one-month, and three-month-old mice via LC–MS/MS. Levels of both EE and DE acylceramides were highest in P0 mice and in one-month-old mice were reduced to 12% (EE) and 23% (DE) of P0 levels. In three-month-old mice they were reduced to 23% (EE) and 50% (DE), respectively, of P0 levels (Fig. 5*C*). Because the levels of EE and DE acylceramides reflect those of the corresponding P-EO (EE) and P-EO (DE) protein-bound ceramides, the marked reduction observed from P0 to adult (one- and three-month-old) mice is consistent with postnatal thinning of the SC. In the dorsal skin, the SC of P0 mice consists of approximately 10 layers with a thickness of about 20 µm (34), whereas that of adult haired mice comprises only 2–3 layers with a thickness of approximately 3 µm (35). This thinning of the SC likely reflects a reduced relative contribution of the SC to the permeability barrier as hair and sebum increasingly contribute to barrier formation during postnatal development. The DE/EE ratio increased from 0.07 in P0 mice to approximately 0.14 and 0.15 in one- and three-month-old mice, respectively, representing an approximate doubling relative to P0 mice (Fig. 5*D*), with no substantial difference between the one- and three-month-old groups. In summary, the DE/EE ratio in mice was higher in the upper layers of the SC and in adult mice than in the epidermis and neonatal mice, respectively. However, even under these conditions, the ratio remained much lower than that observed in human SC (approximately six; Fig. 3*E*). Therefore, the predominance of P-EO (DE) in humans and P-EO (EE) in mice cannot be explained by differences in sampling layer or age, but rather reflects a species-specific difference.

### Involvement of EPHX3 in the stereospecific production of (9*R*,10*S*) DE acylceramides

The conversion of EE acylceramides to DE acylceramides involves the hydrolysis of an epoxide group. Among the epoxide hydrolase family (EPHX1–4), EPHX2 and EPHX3 are candidate enzymes for this reaction, since *in vitro* they hydrolyze a free EH fatty acid, which is used as a model substrate corresponding to the *O*-acyl chain of EH acylceramides, producing a TH fatty acid (Fig. 6*A*) (36). The *O*-acyl chain of epidermal EH acylceramides has been shown to comprise exclusively (9*R*,10*R*,13*R*) EH fatty acids (37). Both EPHX2 and EPHX3 hydrolyze (9*R*,10*R*,13*R*) EH fatty acids to generate (9*R*,10*S*,13*R*) TH fatty acids (Fig. 6*A*), whereas EPHX2 also hydrolyzes (9*R*,10*S*,13*R*) EH fatty acids, reflecting its broader substrate specificity (36). In *Ephx3* KO mice, a slight increase in transepidermal water loss (indicative of a weak skin barrier defect) and an approximately 40% reduction in protein-bound ceramides relative to wild-type mice were observed, whereas *Ephx2* KO mice showed transepidermal water loss comparable to that of wild-type mice (38). To obtain clues as to whether EPHX2 and EPHX3 are involved in the production of P-EO (DE) ceramides, we examined their expression levels in the skin of P0 and one- and three-month-old mice via quantitative real-time RT-PCR. Expression levels of *Ephx3* were 2.6- and 4.2-fold those in P0 mice in one- and three-month-old mice, respectively (Fig. 6*B*). In contrast, expression levels of *Ephx2* were approximately half of P0 levels at one month and had returned to P0 levels by three months.

**Fig. 6.**
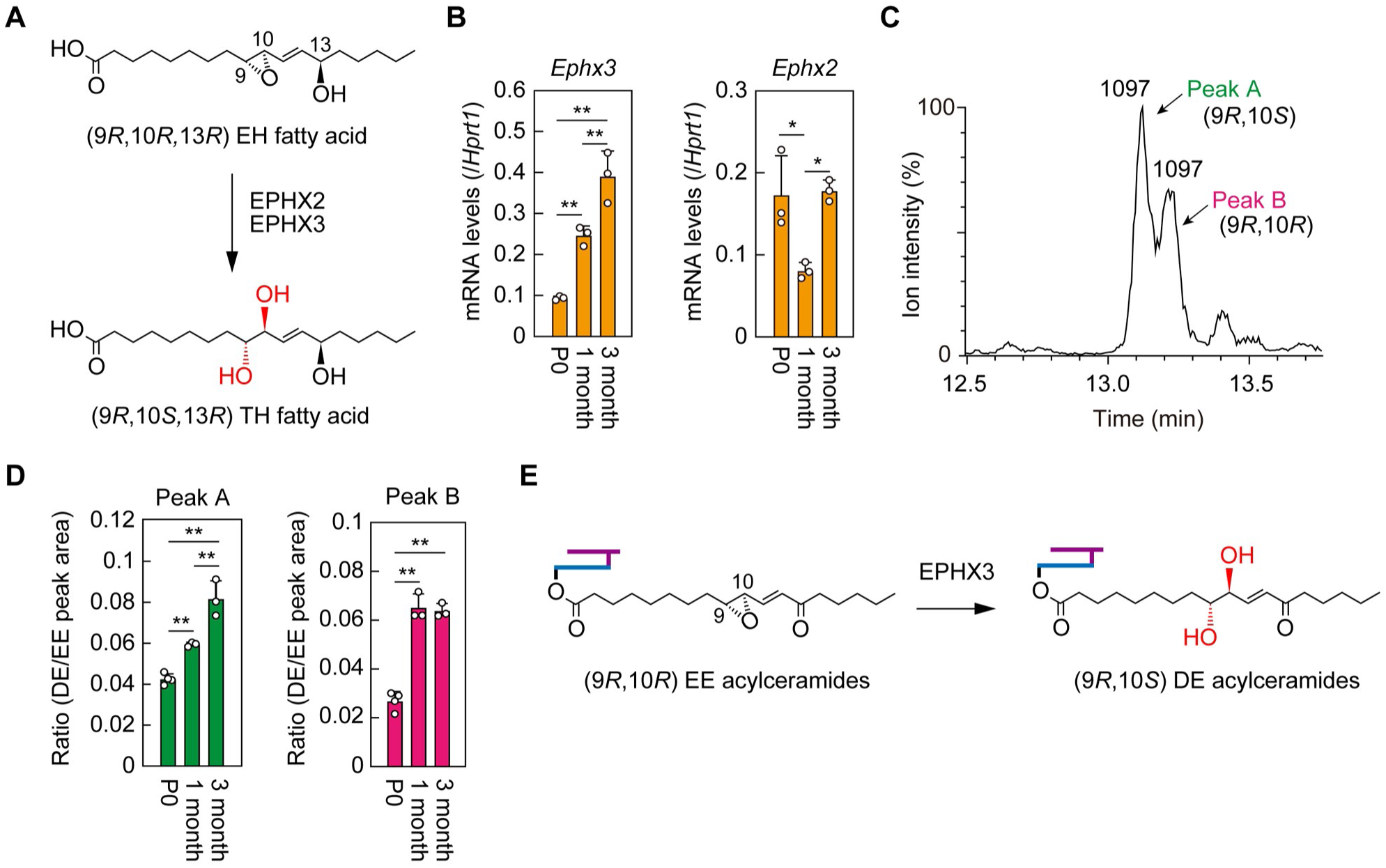
Age-dependent increases in *Ephx3* expression and the (9*R*,10*S*) stereoisomer of P-EO (DE) ceramides in mouse epidermis. (*A*) Schematic representation of the conversion of the (9*R*,10*R*,13*R*) EH fatty acid to the (9*R*,10*S*,13*R*) TH fatty acid mediated by the epoxide hydrolase EPHX2 and EPHX3. (*B*–*D*) Total RNA (*B*) or protein-bound ceramide fractions (*C* and *D*) were prepared from the skin of mice at postnatal day 0 (P0), one month, and three months of age. (*B*) Expression levels of *Ephx2* and *Ephx3* were quantified via quantitative real-time RT–PCR and normalized to the housekeeping gene *Hprt1*. Values presented are means + SD (n = 3; * *P* < 0.05, ** *P* < 0.01; Tukey’s test). (*C* and *D*) DE and EE acylceramides were released from the protein-bound ceramide fraction via oxidative sulfoxide elimination and individually quantified via LC–MS/MS in the MRM mode. (*C*) Representative MRM chromatogram of the DE acylceramide containing C34:1 *N*-acyl chain. Peaks A and B are presumed to correspond to the (9*R*,10*S*) and (9*R*,10*R*) stereoisomers, respectively. (*D*) Values presented are means + SD of the ratios of DE to EE acylceramide peak areas for peaks A and B, respectively (n = 3; * *P* < 0.05, ** *P* < 0.01; Tukey’s test). (*E*) Schematic illustration of a proposed stereospecific cleavage of EE acylceramides into DE acylceramides mediated by EPHX3. In the schematic diagrams, the long-chain base and *N*-acyl chain are shown in purple and blue, respectively.

The specific ions derived from DE acylceramides that were prepared from human SC appeared as paired LC peaks with identical *m/z* values (Fig. 2*A*). These peaks are assumed to represent diastereomers, but their stereochemical configurations remain unknown. It has been reported that alkaline hydrolysis of esterified lipids—most likely derived from TH acylceramides—in the free lipid fraction (extractable with organic solvents) of human and porcine epidermis releases TH fatty acids with the (9*R*,10*S*,13*R*) isomer being the most abundant, followed by the (9*R*,10*R*,13*R*) isomer (37). Therefore, the DE acylceramides observed in human SC are thought to consist of the (9*R*,10*S*) and (9*R*,10*R*) stereoisomers. The (9*R*,10*R*,13*R*) and (9*R*,10*S*,13*R*) isomers of TH fatty acids are eluted in the order (9*R*,10*R*,13*R*) followed by (9*R*,10*S*,13*R*) in normal-phase LC (37). This elution order is expected to be reversed in reversed-phase LC, which was used in the present study. Therefore, DE acylceramide peaks 1 and 3 observed in human SC (Fig. 2*A*), as well as peak A observed in mouse epidermis (Fig. 6*C*), are expected to correspond to the (9*R*,10*S*) stereoisomer, whereas peaks 2 and 4 in human SC (Fig. 2*A*) and peak B in mouse epidermis (Fig. 6*C*) may correspond to the (9*R*,10*R*) stereoisomer. Peaks A and B represent DE acylceramide species containing C34:1 as the *N*-acyl chain, which is the most abundant DE acylceramide species in mouse epidermis (Fig. 3*D*).

Each of the two DE acylceramide peaks was quantified separately in P0 and one- and three-month-old mice. The quantity of peak A increased with postnatal development (Fig. 6*D*). In contrast, peak B increased 2.4-fold relative to P0 at one month and remained at a similar level at three months. Thus, of the two DE acylceramide peaks, peak A showed a pattern similar to that of *Ephx3* expression levels (Fig. 6*B*). Accordingly, EPHX3 is likely to be involved in the production of the (9*R*,10*S*) stereoisomer of DE ceramides. Indeed, *Ephx3*-deficient mouse epidermis shows a marked and selective reduction in the abundance of (9*R*,10*S*,13*R*) TH fatty acids released from esterified epidermal lipids (38). Combined, our results strongly suggest that, in the pathway of protein-bound ceramide production, EPHX3 is involved in the generation of (9*R*,10*S*) DE acylceramides (Fig. 6*E*), leading to the downstream P-EO (DE) protein-bound ceramides.

## Discussion

We previously demonstrated that P-EO (EE) ceramides are the predominant protein-bound ceramides in mice (27). In contrast, the present study reveals that in humans these are minor components, with P-EO (DE) ceramides instead being the dominant protein-bound ceramides (Fig. 3). DE acylceramide ions with specific *m/z* values, released by oxidative sulfoxide elimination of P-EO (DE) ceramides, were detected as two distinct LC peaks (Figs. 2*A* and 6*C*). These peaks were predicted to correspond to the (9*R*,10*S*) and (9*R*,10*R*) stereoisomers. P-EO (DE) ceramides are thought to be generated from the DE acylceramides that are first produced by epoxide ring opening of EE acylceramides, prior to their conversion into the protein-bound form (Fig. 7). Alternatively, DE acylceramides may be generated via a pathway in which EH acylceramides are first converted into TH acylceramides and then oxidized to DE acylceramides (Fig. 7). Enzymes of the epoxide hydrolase (EPHX) family, particularly EPHX3, may participate in this process. The age-dependent expression pattern of *Ephx3* in mice suggests that EPHX3 contributes to the production of (9*R*,10*S*) DE acylceramides (Fig. 6). Generation of the (9*R*,10*R*) stereoisomer, however, may involve EPHX2 or other epoxide-metabolizing enzymes.

**Fig. 7.**
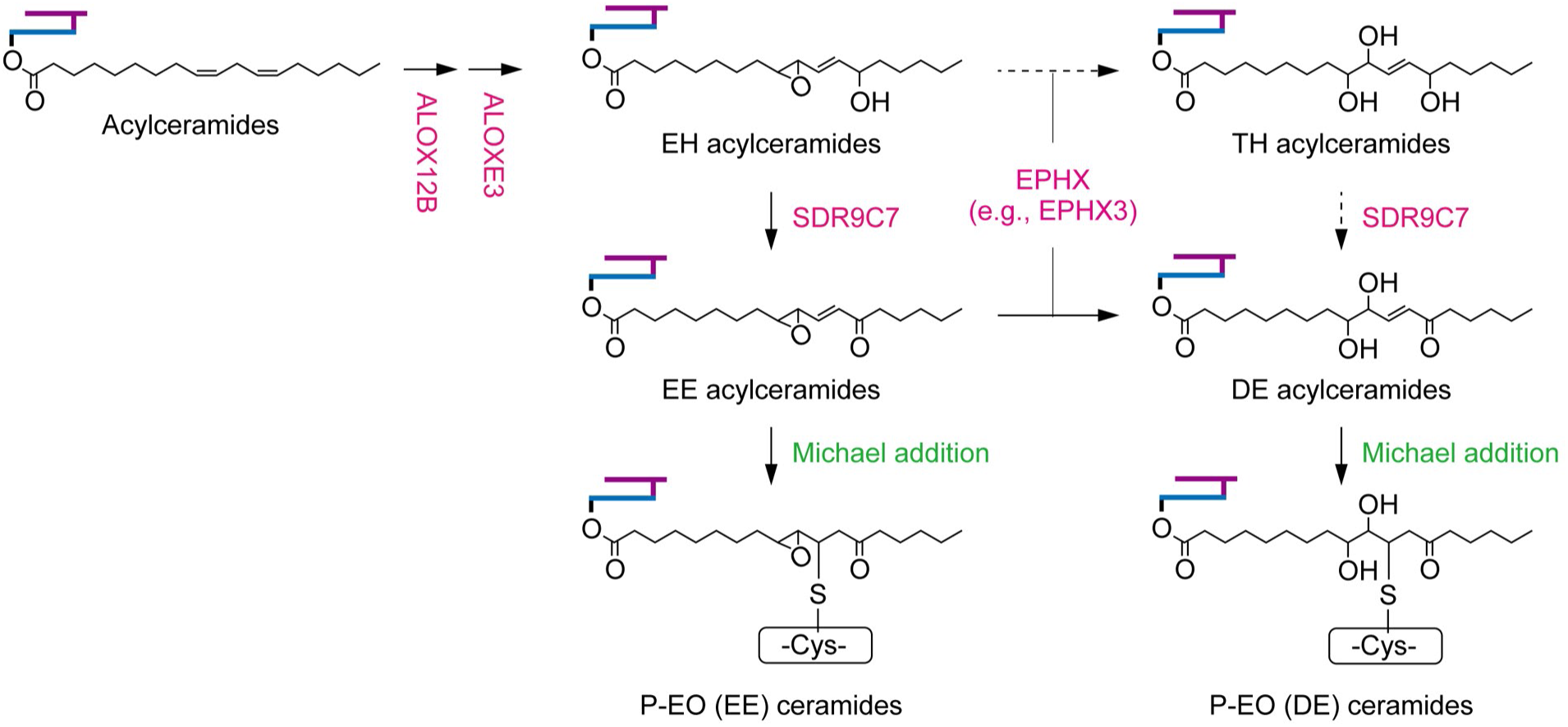
Proposed biosynthetic pathways for P-EO–type protein-bound ceramides. Acylceramides are first converted into EH acylceramides through ALOX12B-dependent hydroperoxidation followed by ALOXE3-mediated hydroperoxide isomerization and are subsequently oxidized by SDR9C7 to yield EE acylceramides. A portion of EH acylceramides can be converted into TH acylceramides via epoxide opening catalyzed by EPHX family enzymes (e.g., EPHX3). Although this conversion is not predominant under physiological conditions, it can become dominant and lead to accumulation in patients with ichthyosis carrying *SDR9C7* mutations (23, 41). Predominantly in mice and partially in humans, EE acylceramides undergo Michael addition to cysteine residues of corneocyte surface proteins to form P-EO (EE) protein-bound ceramides. Alternatively, mainly in humans and partially in mice, EE acylceramides are converted into DE acylceramides by epoxide hydrolase(s) and subsequently form P-EO (DE) protein-bound ceramides via Michael addition. In addition, DE acylceramides may also be generated through a pathway in which EH acylceramides are first converted into TH acylceramides and then oxidized to DE acylceramides; both steps are shown as dashed arrows. In the schematic diagrams, the long-chain base and *N*-acyl chain are shown in purple and blue, respectively.

Our previous analysis using EE and DE analogs revealed that their reactivity toward cysteine residues at the enone moiety was essentially equivalent (27). This raises the question of why the predominant protein-bound ceramide type differs between humans and mice, with DE-type ceramides predominating in humans and EE-type ceramides in mice. Humans and mice differ in the relative importance of the SC in forming the permeability barrier. In adult mice in particular, as hair development progresses, the contribution of hair and sebum to the permeability barrier increases, concomitant with a reduced thickness and contribution of the SC. In contrast to mice, humans never develop extensive body hair, resulting in a thicker SC and a greater contribution of the SC to the permeability barrier. Such differences in the relative contribution of the SC to the permeability barrier may partially explain the distinct ceramide compositions observed in humans and mice. Although sphingosine is the predominant long-chain base in the free ceramides of mouse SC, humans predominantly contain long-chain bases with an additional hydroxyl group, such as 6-hydroxysphingosine and phytosphingosine (31). A similar difference is observed for protein-bound ceramides, as human P-EO (DE) ceramides contain two more hydroxyl groups than mouse P-EO (EE) ceramides. Therefore, free ceramides in the lipid lamellae and protein-bound ceramides forming the CLE are expected to interact more strongly in humans than in mice through hydrogen bonding between these hydroxyl groups, contributing to the formation of a more robust permeability barrier. Thus, protein-bound ceramides may have adapted in humans and mice, respectively, to be optimally matched to the free ceramide composition in each species. Alternatively, the difference in protein-bound ceramides between humans and mice may influence the efficiency of conversion into irreversible forms.

The differences in levels of DE-type protein-bound ceramides between humans and mice may reflect differences in epidermal expression of the epoxide hydrolase genes *EPHX2/3* in humans and *Ephx2/3* in mice. Based on RNA-seq data from the Human Protein Atlas and the Genotype–Tissue Expression databases (https://www.proteinatlas.org/ENSG00000105131-EPHX3/tissue/Skin#rnaseq, https://www.proteinatlas.org/ENSG00000120915-EPHX2/tissue/Skin#rnaseq, https://www.proteinatlas.org/ENSG00000165704-HPRT1/tissue/Skin#rnaseq), the expression levels of *EPHX2* and *EPHX3* in human skin were estimated to be 2–3-fold and 4–6-fold, respectively, the transcripts per million (TPM) value of *HPRT1*, a housekeeping gene. In contrast, analysis of skin samples from three-month-old mice via quantitative real-time RT-PCR in this study revealed *Ephx2* and *Ephx3* expression levels of 0.18 and 0.40, respectively, of *Hprt1* levels (Fig. 6*B*). Consistently with this, RNA-seq data from the Expression Atlas (https://www.ebi.ac.uk/gxa/home) indicated TPM ratios of *Ephx2* and *Ephx3* to *Hprt1* of approximately 0.44 and 0.58, respectively. Combined, these findings indicate that although *Ephx2* and *Ephx3* are expressed in mouse skin, their expression levels are lower than those of the corresponding *EPHX* genes in human skin.

Although P-EO (EE) and P-EO (DE) ceramides can be reversibly released into organic solvents via oxidative sulfoxide elimination, approximately 40% of the protein-bound ceramides in both human SC (Fig. 4) and mouse epidermis (27) are irreversibly bound. The structures of these irreversibly bound protein-bound ceramides have not yet been elucidated. From a structural standpoint, P-O–type protein-bound ceramides are classified within this irreversibly bound form (Fig. 4*A*). The P-O–type ceramide has historically been proposed as a structural model of protein-bound ceramides; however, its existence has not been confirmed by definitive experimental evidence. The P-O–type ceramide model was proposed based on the following observations. In 1987, Wertz and Downing found that mild alkaline treatment of lipid-depleted residues prepared from porcine SC resulted in the release of ω-hydroxyceramides, providing evidence for the existence of covalently bound ceramides (39). Swartzendruber and colleagues subsequently demonstrated that extraction of free lipids from isolated SC with chloroform/methanol resulted in the disappearance of the lipid lamellae, while the CLE was preserved; this structure was then removed by alkaline treatment (26). These observations suggested an ester linkage as the mode of attachment, given the alkaline sensitivity of the bound ceramides. Furthermore, the high content of glutamate and aspartate residues in cornified envelope proteins, together with considerations from molecular modeling, prompted the proposal of the P-O model, in which ω-hydroxyceramides are ester-linked to protein carboxyl groups (26). In 1998, Marekov and Steinert reported that digestion of a cornified envelope protein fraction derived from human foreskin with Proteinase K yielded peptides covalently linked to ceramides, all of which contained glutamate residues (40). Due to the analytical difficulty of this type of investigation, this remains the only study that has analyzed protein-bound ceramides in a peptide-linked state. Although that study did not explicitly classify the structural type of the protein-bound ceramides, comparison of the masses of the peptide–protein-bound ceramide conjugates before and after alkaline treatment is currently interpreted as being more consistent with P-O–type protein-bound ceramides than with the P-EO–type proposed later. In this context, the 1998 Marekov and Steinert study (40) may be the only experimental report that provides support for the P-O–type ceramide hypothesis.

In our previous study, the protein-bound ceramide fraction isolated from mouse epidermis was treated with Pronase E to digest the protein moiety of the protein-bound ceramides down to the amino acid level, followed by LC–MS/MS analysis using a precursor-ion scan targeting sphingosine-containing ceramides. P-EO (EE) protein-bound ceramides were detected, but no P-O–type protein-bound ceramides were observed (27). These results suggest that, at least in mouse epidermis, P-O–type protein-bound ceramides are not a major component of the total pool of protein-bound ceramides, but instead may exist as a limited subset or be detectable only under specific experimental conditions. Indeed, in the study by Marekov and Steinert (40), cornified envelope fractions were subjected to mild saponification, and the possibility that this treatment may have secondarily generated structures resembling P-O–type ceramides cannot be excluded. We consider it likely that the majority of irreversibly bound protein-bound ceramides are not of the P-O type but are instead metabolites of or structurally related to P-EO (EE) or P-EO (DE) ceramides. One possible scenario is that EE or DE acylceramides initially form a Schiff base with an amino acid residue of a corneocyte surface protein. The resulting imine intermediate may then be attacked by a thiol group, derived either intramolecularly from the same protein or intermolecularly from another protein, leading to thioamination at the imine carbon.

In this study, we not only identified P-EO (DE) ceramides as protein-bound ceramides in the human SC for the first time, but also demonstrated that they represent the predominant type. Protein-bound ceramides are essential for epidermal barrier function, and defects in their production lead to ichthyosis (12, 13). Therefore, the findings of the present study contribute to a deeper understanding of the molecular mechanisms underlying skin barrier formation and the pathology thereof. Future studies clarifying the involvement of EPEX2/3 in P-EO (DE) ceramide production are expected to elucidate the complete biosynthetic pathway of protein-bound ceramides. Furthermore, determination of the structures of irreversibly bound forms of protein-bound ceramides should reveal the full spectrum of protein-bound ceramide types present in the human SC.

## Materials and Methods

### Collection of human SC by tape stripping

Human SC samples were collected from healthy volunteers (n = 6; males and females, aged 20–40 years) after obtaining written informed consent from all participants. The inner forearm was gently cleaned with water and air-dried, and SC samples were collected by applying and removing film masking tape (No. 465#40; Teraoka Seisakusho, Tokyo, Japan). This procedure was repeated three times at the same site. The second tape strip was used for lipid extraction because the first strip may have been contaminated with dust, detergents, or lotions. This study was approved by the Ethics Committee of Hokkaido University (approval no. 2018-004) and was conducted in accordance with the Declaration of Helsinki and the relevant institutional guidelines.

### Mice

Male and female C57BL/6J mice (Japan SLC, Shizuoka, Japan) were used in this study. The animals were maintained under specific pathogen-free conditions at 23 ± 1 °C with 50 ± 10% humidity and a 12 h light/dark cycle, with free access to water and standard chow (CLEA Rodent Diet CE-2; CLEA Japan, Tokyo, Japan). All animal experiments were approved by the Animal Care and Use Committee of Hokkaido University (approval no. 22-0036) and were conducted in accordance with institutional guidelines.

### Collection of mouse epidermis and SC samples

To isolate the epidermis, skin from P0 mice was removed from the whole body after sacrifice by decapitation. For one- and three-month-old mice, the dorsal skin was harvested after shaving followed by hair removal with forceps. The skin was incubated with 2 mg/mL Dispase II (Thermo Fisher Scientific, Waltham, MA, USA) in phosphate-buffered saline at 4 °C for 16 h. The epidermis was separated from the dermis using forceps, washed with phosphate-buffered saline, pressed onto filter paper to remove excess water, and weighed. Mouse SC samples were collected from P0 mice by tape stripping over the entire body surface using the same procedure as for human SC sampling.

### Oxidative sulfoxide elimination-mediated release of DE and EE acylceramides from protein-bound ceramides

Human and mouse SC were recovered from tape strips by ultrasonication in 1 mL of methanol for 5 min at room temperature, followed by removal of the tape. To prepare mouse epidermal samples, the epidermis (5 mg) was transferred into a tube containing zirconia beads (1 mm in diameter; TOMY Seiko, Tokyo, Japan), suspended in 500 µL CHCl_3_/CH_3_OH (1:2, v/v), and homogenized (4,500 rpm, 4 °C, 1 min) using a Micro Smash MS-100 (TOMY Seiko). Human and mouse SC samples, as well as mouse epidermal samples, were then suspended in 2 mL of CHCl_3_/CH_3_OH (1:2, v/v), vigorously mixed, centrifuged (2,600 × *g*, room temperature, 3 min), and the supernatant was discarded. This procedure was repeated five times to completely remove free lipids. The resulting pellet (protein-bound ceramide fraction) was collected, dried, and weighed. The pellet was suspended in CHCl_3_/CH_3_OH (1:2, v/v) at 1 mg/mL and aliquots of 20 µL (0.02 mg, human SC) or 50 µL (0.05 mg, mouse epidermis/SC) were transferred to new tubes.

DE and EE acylceramides were released from the protein-bound ceramide fraction by five successive rounds of oxidative sulfoxide elimination, essentially as described previously (27, 28). Briefly, the protein-bound ceramide fraction was suspended in 600 µL of CHCl_3_/CH_3_OH (1:2, v/v) and incubated at 60 °C for 3 h (the first, second, fourth, and fifth rounds) or 15 h (the third round), followed by centrifugation (20,400 × *g*, room temperature, 3 min). The supernatant was collected and 600 µL of CHCl_3_/CH_3_OH (1:2, v/v) was added to the pellet in each round. The supernatants from all rounds were combined, dried, and suspended in 375 µL of CHCl_3_. As an internal standard, 1 pmol of nine deuterium (*d*_9_)-labeled acylceramide (*N*-[26-oleoyloxy(*d*_9_) hexacosanoyl]-D-*erythro*-sphingosine; Avanti Polar Lipids, Alabaster, AL, USA) was added, and the mixture was then combined with 375 µL of CH_3_OH and 337.5 µL of H_2_O. After centrifugation (20,400 × *g*, room temperature, 3 min), the lower (organic) phase was collected, dried, dissolved in 300 µL of CH_3_OH, and subjected to LC–MS/MS analysis as described below (injection volume, 5 µL).

### Preparation and quantification of reversibly and irreversibly bound protein-bound ceramides

The protein-bound ceramide fraction (0.02 mg) prepared from human SC was subjected to six successive rounds of oxidative sulfoxide elimination by incubation in 600 µL of CHCl_3_/CH_3_OH (1:2, v/v) at 60 °C (3 h for the first, second, fourth, fifth, and sixth rounds; 15 h for the third round), followed by centrifugation (20,400 × *g*, room temperature, 3 min) after each round. The supernatants from all rounds were collected, dried, and designated as the reversible protein-bound ceramide fraction. The pellet remaining after the six extractions was used as the irreversible protein-bound ceramide fraction. The protein-bound ceramide fraction before oxidative sulfoxide elimination (total fraction), as well as the reversibly and irreversibly bound protein-bound ceramide fractions, were mixed with 400 µL of 1 M KOH in 95% CH_3_OH containing 1 pmol of *N*-(2’-(*R*)-hydroxypalmitoyl(*d_9_*))-D-*erythro*-sphingosine (*d*_9_-C16:0 α-hydroxyceramide; Avanti Polar Lipids) as an internal standard and incubated at 60 °C for 1 h to hydrolyze the ester linkages and release ω-hydroxyceramides from each fraction. The samples were neutralized by adding 22.9 µL of 17.5 M acetic acid and then combined with 380 µL of CHCl_3_ and 300 µL of H_2_O. After centrifugation (20,400 × *g*, room temperature, 3 min), the lower (organic) phase was collected, dried, dissolved in 1 mL of CH_3_OH, and subjected to LC–MS/MS analysis (injection volume, 5 µL) to quantify ω-hydroxyceramides as described below.

### Ceramide analysis via LC–MS/MS

Ceramides were analyzed using ultra-performance LC coupled with a tandem triple quadrupole mass spectrometer (Xevo TQ-XS; Waters, Milford, MA, USA) as described previously (27). LC separation was performed using a reversed-phase column (ACQUITY UPLC CSH C18 column; particle size, 1.7 µm; inner diameter, 2.1 mm; length, 100 mm; Waters) with mobile phase A (CH_3_CN/H_2_O [3:2, v/v] containing 5 mM ammonium formate) and mobile phase B (CH_3_CN/2-propanol [1:9, v/v] containing 5 mM ammonium formate), as described previously (27). Lipids were ionized via electrospray ionization in positive or negative ion modes under the following conditions: capillary voltage, 2.5 kV; source offset, 50 V; desolvation temperature, 650 °C; desolvation gas flow, 1,200 L/h; cone gas flow, 150 L/h; nebulizer gas pressure, 7.0 bar. The cone voltage was set to 30 V for precursor ion scanning and 35 V for product ion scanning in the positive ion mode, and to 30 V for product ion scanning in the negative ion mode. The collision energy was set to 40 eV for precursor ion scanning and to 30 eV for product ion scanning in both positive and negative ion modes. MRM was performed for the detection of EE acylceramides, DE acylceramides, and ω-hydroxyceramides using the mass-spectrometric parameters listed in Table 1.

**Table 1.**
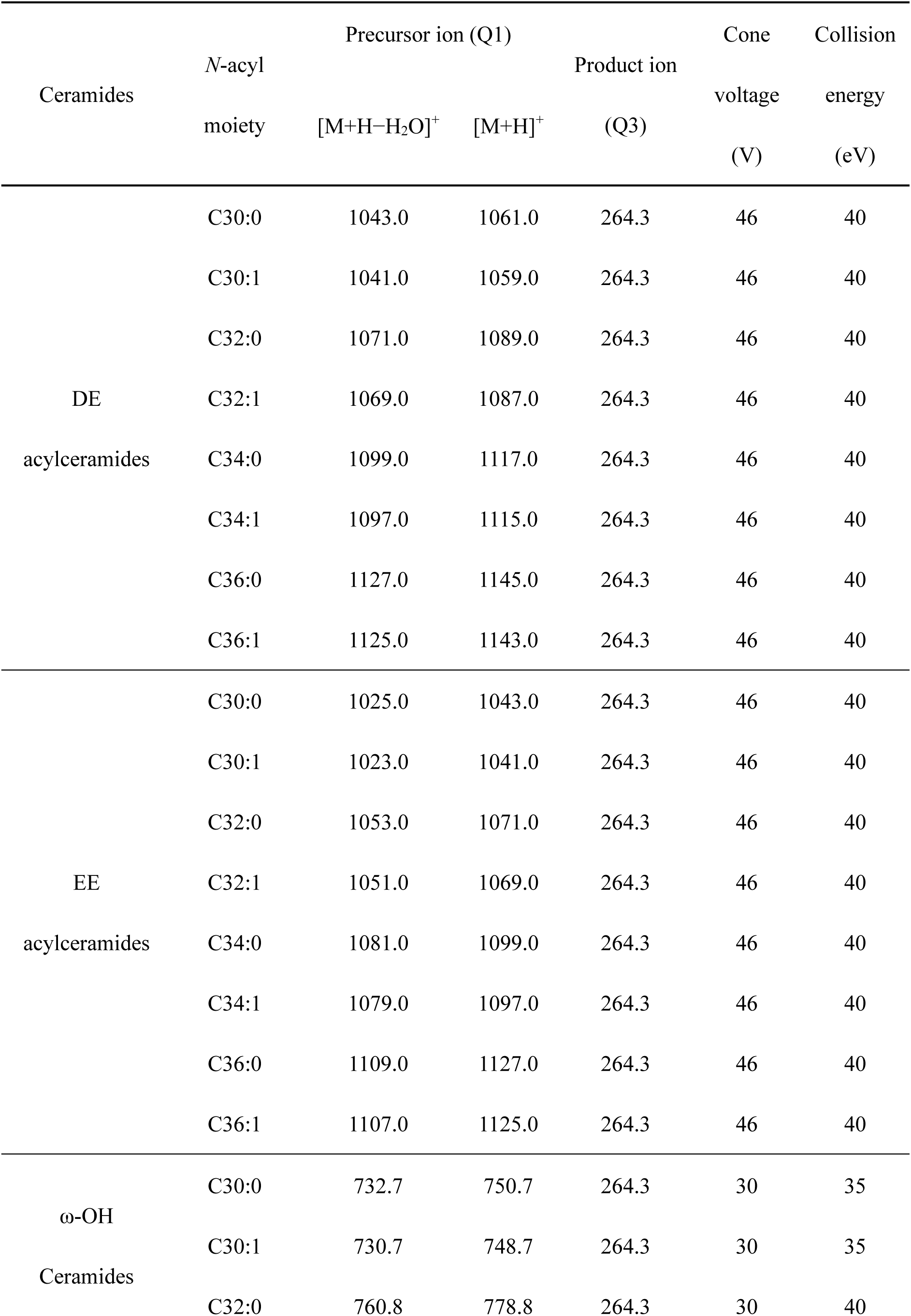
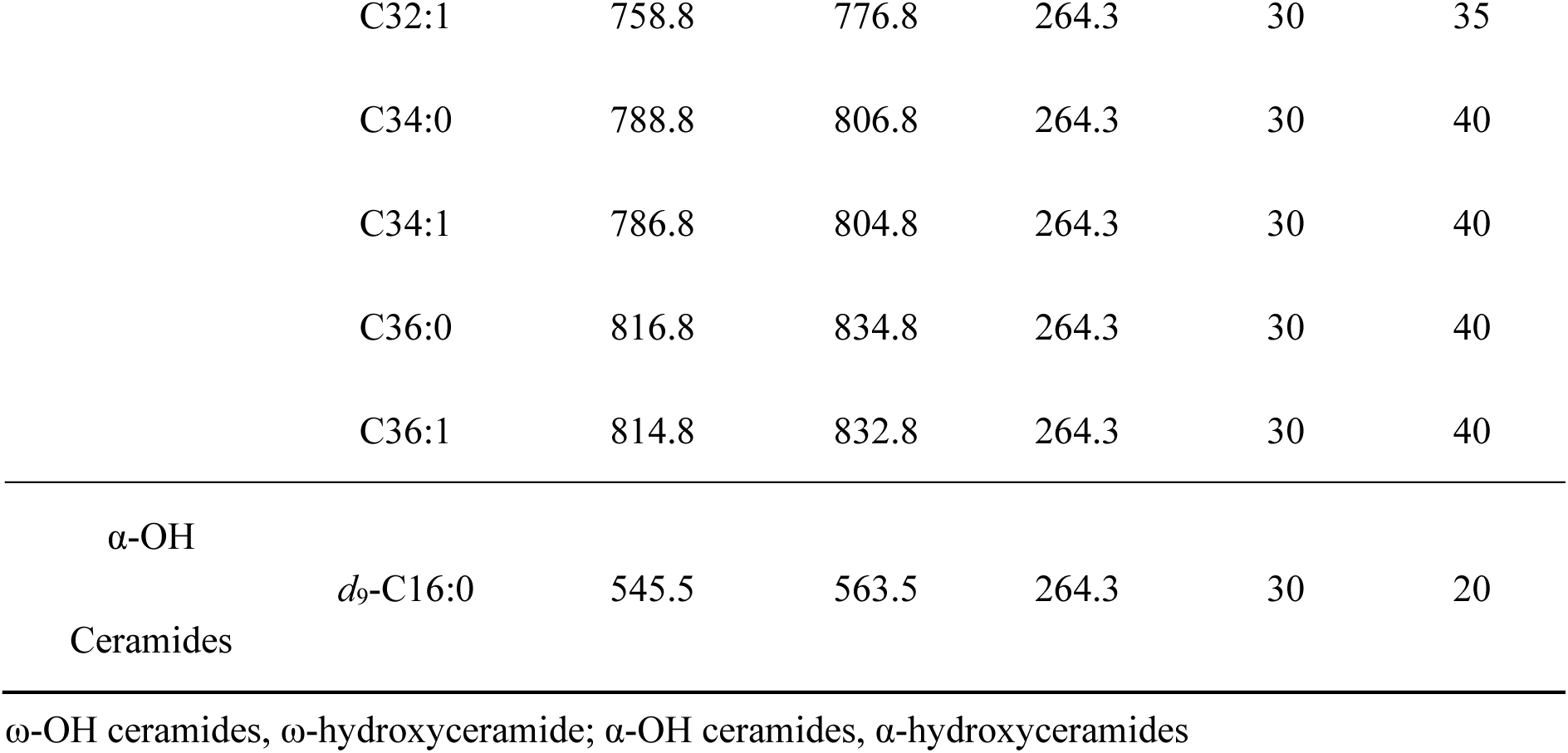
MRM parameters for detection of ceramide species in LC–MS/MS analyses.

| Ceramides | *N*-acyl moiety | Precursor ion (Q1) |  | Product ion (Q3) | Cone voltage (V) | Collision energy (eV) |
| --- | --- | --- | --- | --- | --- | --- |
|  |  | [M+H–H<sub>2</sub>O]<sup>+</sup> | [M+H]<sup>+</sup> |  |  |  |
| DE<br>acylceramides | C30:0 | 1043.0 | 1061.0 | 264.3 | 46 | 40 |
|  | C30:1 | 1041.0 | 1059.0 | 264.3 | 46 | 40 |
|  | C32:0 | 1071.0 | 1089.0 | 264.3 | 46 | 40 |
|  | C32:1 | 1069.0 | 1087.0 | 264.3 | 46 | 40 |
|  | C34:0 | 1099.0 | 1117.0 | 264.3 | 46 | 40 |
|  | C34:1 | 1097.0 | 1115.0 | 264.3 | 46 | 40 |
|  | C36:0 | 1127.0 | 1145.0 | 264.3 | 46 | 40 |
|  | C36:1 | 1125.0 | 1143.0 | 264.3 | 46 | 40 |
| EE<br>acylceramides | C30:0 | 1025.0 | 1043.0 | 264.3 | 46 | 40 |
|  | C30:1 | 1023.0 | 1041.0 | 264.3 | 46 | 40 |
|  | C32:0 | 1053.0 | 1071.0 | 264.3 | 46 | 40 |
|  | C32:1 | 1051.0 | 1069.0 | 264.3 | 46 | 40 |
|  | C34:0 | 1081.0 | 1099.0 | 264.3 | 46 | 40 |
|  | C34:1 | 1079.0 | 1097.0 | 264.3 | 46 | 40 |
|  | C36:0 | 1109.0 | 1127.0 | 264.3 | 46 | 40 |
|  | C36:1 | 1107.0 | 1125.0 | 264.3 | 46 | 40 |
| ω-OH<br>Ceramides | C30:0 | 732.7 | 750.7 | 264.3 | 30 | 35 |
|  | C30:1 | 730.7 | 748.7 | 264.3 | 30 | 35 |
|  | C32:0 | 760.8 | 778.8 | 264.3 | 30 | 40 |

### Quantitative real-time RT-PCR

Mouse skin was collected as described above, immersed in RNAlater (Merck, Darmstadt, Germany), cut into small pieces, and transferred to a tube containing zirconia beads. After the addition of 1 mL of TRIzol Reagent (Thermo Fisher Scientific), the skin was homogenized using a Micro Smash MS-100 (5,000 rpm, 4 °C, 1 min). This homogenization procedure was repeated five times at 1 min intervals, with the samples chilled on ice between repetitions. After the addition of 200 µL of CHCl_3_, the samples were vigorously mixed for 3 min and centrifuged (12,000 × *g*, 4 °C, 15 min). The upper aqueous phase was collected, mixed with 500 µL of isopropanol, and then incubated at room temperature for 10 min to precipitate RNAs. After centrifugation (12,000 × *g*, 4 °C, 10 min), the resulting pellet was washed with 75% ethanol, dried, and dissolved in RNase-free water by incubation at 55 °C for 10 min. The total RNA obtained was converted to cDNA using PrimeScript RT Master Mix (Perfect Real Time; Takara, Shiga, Japan) and oligo(dT) primer. Quantitative real-time PCR was performed using cDNA as a template with KOD SYBR qPCR Mix (TOYOBO, Osaka, Japan) and gene-specific primers (*Ephx2*, 5′-GAATATGCCATGGAATTGCTGTGTA-3′ and 5′-AGATCGGATAACTTTCATGGGAGAC-3′; *Ephx3*, 5′-TCCAAGAATACTCAATCCACCACAT-3′ and 5′-AAAGGAATGCTTCAAGTTCAGAAGG-3′; *Hprt1*, 5′-GCTGACCTGCTGGATTACATTAAAG-3′ and 5′-CTTAACCATTTTGGGGCTGTACTGC-3′) on CFX96 Touch Real-Time PCR Detection System (Bio-Rad, Hercules, CA, USA). The expression levels of *Ephx2* and *Ephx3* were calculated relative to *Hprt1* (hypoxanthine-guanine phosphoribosyltransferase 1).

### Statistical analyses

Data are presented as means + SD. Statistical analyses were performed using two-tailed Welch’s *t*-test, two-tailed Tukey’s test, and Dunnett’s test (Prism; Dotmatics, Boston, MA, USA). A *P*-value of < 0.05 was considered statistically significant.

## ACKNOWLEDGEMENTS

We thank Dr. Satoshi Ichikawa (Hokkaido University), Dr. Akira Katsuyama (Hokkaido University), and Dr. Hiroshi Tanaka (Juntendo University) for insightful discussions. This work was supported by funding from the Takeda Science Foundation (to A.Kihara) and by Japan Society for the Promotion of Science (JSPS) KAKENHI grant numbers JP22H04986 (to A.Kihara) and JP25K24525 (to A.Kihara).

